# Effect of cultivation system on quality changes in durum wheat grain and flour produced in North-Eastern Europe

**DOI:** 10.1101/2020.07.13.200410

**Authors:** Joanna K. Banach, Katarzyna Majewska, Krystyna Żuk-Gołaszewska

## Abstract

Grain of the highest hardness was produced from durum wheat grown without the growth regulator, at the lowest sowing density (350 seeds m^-2^) and nitrogen fertilization dose of 80 kg ha^-1^. The highest values L* and b* were determined in the grain of wheat cultivated without additional agrotechnical measures (growth regulator and nitrogen fertilization). Study results, supported by correlation analysis, indicated that high-quality grain with desired flour quality parameters (extraction rate, granulation and lightness) can be produced from spring durum wheat grown without the growth regulator and at medium doses of nitrogen fertilization. Additionally, this variant of applied cultivation system can reduce costs of durum wheat production and contamination of the natural environment.

## 1. Introduction

Next to maize and rice, wheat is one of the key cereals grown for consumption purposes (Zarroug et al. 2015). Among its multiple species and subspecies, the most economically significant ones include: common wheat (*Triticum aestivum*) representing soft wheats, and durum wheat (*Triticum durum)* (De Santis et al. 2017; 2018). Considering its hardness and vitreousness as well as a high content of carotenoid pigments, the grain of *Triticum durum* is mainly used to produce special coarse flour (semolina), which in turn is used for making high-quality pasta, like e.g. spaghetti and penne (Su, He and Sun 2015; Miskelly 2017; Hare 2017). In addition, due to high contents of protein (14 – 15.5%) and gluten (42 – 46%), as well as a low activity of amylolytic enzymes (falling number: 259 – 358 s), and relatively high contents of vitamins and minerals, it can serve as a raw material to produce functional foods. Flour made of durum wheat grain is used to produce bread loaves or as an additive improving its properties, as well as to produce cake dough, pizza, cookies, couscous, and extruded products (extruded grain) (De Santis et al. 2017).

The technological value of durum wheat grain is principally indicated by its usability to manufacture the final product of a sufficient quality. The main ingredient of premium quality pasta is 100 percent durum wheat semolina. Semolina, which originates from the endosperm of durum wheat grain is essentially coarse flour. Good quality pasta can also be made of durum wheat flour or a blend of semolina and durum wheat flour (granular). Pasta can be produced using common wheat flour, but it is generally inferior in appearance (color). Semolina particle size distribution should be as narrow as possible to minimize uneven hydration during mixing in the pasta making process (Owens 2001). Apart from the pursuit of the best possible financial results by every link of the grain supply chain (farms) and grain processing into the flour (mills) grain hardness and color as well as the quantity and quality of flour made of it are becoming increasingly important (Delcour and Hoseney 2010; Pauly et al. 2013).

Hruškova and Švec (2009) showed that wheat grain hardness results from adhesion between starch granules and storage proteins. Low molecular proteins of starch granules are present in a larger amount in soft wheat compared to hard cultivars. Moreover, non-uniform particle size distribution of semolina results in the uneven hydration and pasta defects (Morris and Fuerst 2015). Grain hardness is also affected by puroindolines, which act to soften the endosperm, but are lacking in durum wheat (Pasha, Anjum and Morris 2010; Boehm et al. 2018). Nagamine et al. (2003) demonstrated a significant correlation between flour color and hardness. In turn, Hruškova and Švec (2009) who analyzed correlations between grain hardness of wheat and its relation to other quality features, showed that the strongest relation was obtained for the grain ash content, semolina yield, and flour protein content.

The yellow-amber color of durum wheat grain, usually associated by consumers with its high quality, is mainly due to the accumulation of two groups of natural pigments: carotenoids and anthocyanins. Carotenoids contribute to the yellow color of the endosperm of durum wheat grain and semolina, whereas anthocyanins accumulate in the pericarp of durum wheat and contribute to the blue, violet, and red color of its grain. Apart from ensuring esthetic values, the pigments contained in the grain play an important nutritional and health role, therefore their modification in the grain by means of agrotechnical factors is still valid and justified in research (Abdel-Aal et al. 2007, Nishino et al. 2009, Beleggia et al. 2010).

Changes in the quality parameters of wheat grain are affected, most of all, by varietal traits, habitat conditions, cultivation system, and agrotechnical measures, including nitrogen fertilization (Makowska et al. 2008; Giuliani et al. 2011; Ficco et al. 2014; Semenov et al. 2014; De Santis et al. 2017; Zuk-Golaszewska et al. 2018). Nitrogen is the major component of fertilizers which significantly influences crop yield and protein content (Giuliani et al. 2011).

Due to the above, the basic task of the modern plant production is to strive for high, stable and good-quality crops, with the lowest possible inputs and respect for the natural environment (Diacono et al. 2013, Ropelewska et al. 2019). Durum wheat is an agronomically competitive crop to common wheat, which exhibits tolerance to biotic and abiotic stress and is widely cultivated in regions with low rainfall (Marti and Slafer 2014; Kaur et al. 2015; Campiglia et al. 2015). Generally, nitrogen fertilizer management may influence wheat grain hardness (Giuliani et al. 2011). Makowska et al. (2008) showed that nitrogen fertilization had a distinct effect on grain hardness and vitreousness, but no significant effect on carotenoids content and color of pasta dough.

Investigations addressing the coupled use of a few treatments in crop cultivation systems, especially in an integrated farming system (Diacono et al. 2013; Walia et al. 2019), differing in nitrogen fertilization level and sowing density are sparse. The growing scale of production and the increasing diversity of wheat applications urge the need for assessing the quality of grain batches received by warehouses. In turn, the production of high-quality flour (semolina) requires sufficient raw material and continuous control of the production process (Mefleh et al. 2018).

Considering the above, a study was undertaken with the aim to determine the effect of applied cultivation system of spring durum wheat: differing in sowing density, retardant administration, and nitrogen fertilization dose, on the selected quality parameters of grain and flour made of it. Moreover, an analysis of correlations between agrotechnical factors and technological parameters was performed to determine which factors significantly shape the technological quality of studied material.

## 2. Material and methods

### 2.1. Cultivation of durum wheat

Study material included grain of spring durum wheat cv. SMH 87 (Polish National List of the Research Center for Cultivar Testing, 2011) originating from the field experiment at the Production and Experimental Station in Bałcyny (53°40’ N, 19°50’E) - harvest 2016, under integrated farming management. This is the first cultivar of durum wheat grown in Poland that is utterly adjusted to its climatic conditions. The experimental variables included:

- nitrogen fertilization: control, without nitrogen fertilization; 80 kg·ha^-1^-50 kg·ha^-1^ before sowing and 30 kg·ha^-1^ at the shooting stage (Z33); 120 kg·ha^-1^-50 kg ha^-1^ before sowing, 30 kg·ha^-1^ at the shooting stage (Z33), and 40 kg·ha^-1^ at the early earing stage (Z51) (Zadoks, Chang, Konzak 1974),
- sowing density: 350; 450, and 550 seeds·m^-2^,
- application of growth regulator: without of application (WGR) or with the application (GR) Medax 350 S.C. as aprophylactic measure that may prevent crop lodging.

After the harvest, the grain was cleansed from physical impurities and residues of the seed coat. It had a standardized moisture content of 14±0.5%, that was assayed acc. to the Polish Standard (2012). The grain (ca. 500 g from each cultivation variant) was poured into glass bottles with a ground-glass stopper and stored in the climatic chamber (ICP 500, Memmert, USA) at air temperature of 20±0.1°C until analyzed.

### 2.1. Measurements grain hardness

Grain hardness was measured with a Universal Testing Device (Instron 5942; Instron Corp., USA). A uniaxial compression test (flat shank: Ø=12.6 mm; speed of the working head in the test: 10 mm/min) was carried outon individual wheat kernels placed on a measuring table crease down, in 20 replications for each sample. Grain hardness was determined based on the measured values of the maximal compression force (F_max_) needed to induce 50% deformation; values were expressed in N. The results of the tests were analyzed using the computer software Bluehill II.

### 2.2. Measurements of grain and flour color

Color measurements were made using a Hunter Miniscan XE Plus colorimeter (HunterLab, USA), which was set to collect spectral data with illuminant A/observer D65/10°. Prior to the measurements, the colorimeter was calibrated using a white and black ceramic plate. Grain and flour color was established based on measurements in the CIELab scale (L*-lightness, a*-redness, b*-yellowness). Color measurements of both grain and flour were conducted in a black container (ca. 50 cm^3^) in 12 replications.

### 2.3. Determination of flour extraction rate

The flour extraction rate (FER) was determined by milling the grain in a laboratory Quadrumat Junior roller mill (Brabender^®^). Before milling, the grain was conditioned to achieve 14.5% moisture content. FER determination was conducted in 6 replications.

### 2.4. Determination of flour particle size (granulation < 400 μm)

Particle size of flour (FPS) obtained from the studied grain was determined according to the method of Haber and Horubałowa (1992). The flour (100 g) was sifted through a sieve with mesh size of 400 μm, using a laboratory sifter (type LPzE-2e, Multiserw Morek, Poland). Percentage contents of flour particle fractions less than 400 μm were determined based on 6 replications.

### 2.5. Statistical analysis

Calculations were carried out in Statistica 13.1 software (StatSoft Inc., Tulsa, OK, USA). The results of analyses were presented as means ± standard deviations. One-way analysis of variance was conducted to determine the influence of agrotechnical factors on the quality parameters of durum wheat grain and flour; its results were reported at the following levels of significance p<0.01 and p<0.05. The relationships between agrotechnical factors and selected quality parameters of grain and flour obtained from it were determined by correlation analysis by calculating the Pearson coefficient at the significance level of p<0.01 and p<0.05 (SPSS ver. 25). However, in order to show statistically significant differences between groups in terms of the use of GR or WGR, the Mann-Whitney U test (nonparametric equivalents of analysis of variance) was conducted. This test allowed verification of the null hypothesis, which assumes that the means of compared groups (seeding density, nitrogen fertilization) show equality and are characterized by a similar distribution.

## 3. Results and discussion

### 3.1. Grain hardness

Increasing the fertilization dose (0, 80, 120 kg·ha^-1^) in the cropping system without the growth regulator (WGR) and at sowing density of 350 and 450 seeds·m^-2^ caused a significant (p<0.01, p<0.05) decrease in the value of the maximal compression force (F_max_) from approx. 145 to 115N and from approx. 135 to 108N, respectively. An opposite observation was made at the higher sowing density (550 seeds·m^-2^), i.e. F_max_ values increased from approx. 116 to approx. 148 N along with an increasing fertilization dose. The statistical analysis of results demonstrated also that the increased nitrogen doses and sowing densities caused significant (p<0.01; p<0.05) differences in F_max_ value determined for the grain from wheat grown in the WGR variant (table 1).

**Table 1.** Changes in the maximum compression force (mean values ± standard deviation) of the durum wheat grain depending on sowing density, growth regulator application and nitrogen fertilization.

| Sowing density<br>(seeds·m<sup>-2</sup>) |  | Maximum compression force - F<sub>max</sub> (N) |  |  |  |  |  |
| --- | --- | --- | --- | --- | --- | --- | --- |
|  |  | Nitrogen fertilization (kg·ha<sup>-1</sup>) |  |  | Significant differences – 350/450/550 |  |  |
|  |  | 0 | 80 | 120 | 0 | 80 | 120 |
| Without the application growth regulator - WGR |  |  |  |  |  |  |  |
| 350 |  | 144.82±35.91<sup>aC</sup> | 139.80±6.76<sup>a</sup> | 114.42±11.51<sup>bB</sup> | * | ** | ** |
| 450 |  | 135.11±8.90<sup>Ab</sup> | 119.44±17.64<sup>Ba</sup> | 108.11±15.33<sup>a</sup> |  |  |  |
| 550 |  | 115.73±9.31<sup>a</sup> | 117.24±10.42<sup>a</sup> | 147.92±23.86<sup>b</sup> |  |  |  |
| With the application growth regulator - GR |  |  |  |  |  |  |  |
| 350 |  | 116.17±16.28<sup>a</sup> | 114.52±15.32<sup>aA</sup> | 137.20±25.13<sup>Ba</sup> | NS | NS | NS |
| 450 |  | 118.17±10.55<sup>Ac</sup> | 108.73±6.27<sup>Bb</sup> | 127.70±10.66<sup>c</sup> |  |  |  |
| 550 |  | 128.80±15.12<sup>a</sup> | 134.57±20.62<sup>a</sup> | 136.12±17.45<sup>a</sup> |  |  |  |
| Significant<br>differences –<br>GR/WGR | 350 | NS | ** | * | - | - | - |
|  | 450 | ** | NS | ** |  |  |  |
|  | 550 | NS | NS | NS |  |  |  |

\* – significant differences, p<0.05; \*\* – significant differences, p<0.01; NS – no significant differences
a-d – mean values in columns indicated with various small letters are statistically significantly different at p<0.01;
A-B – mean values in columns indicated with various capital letter are statistically significantly different at p<0.05
a-a – mean values in columns indicated with the same small letters non differ statistically

A strong impact of fertilization with nitrogen on the increased hardness of the grain was also reported by other authors (Ciołek and Makarska 2004; Makowska et al. 2008; Szumilo and Rachon 2010; Rachon et al. 2015). So explicit tendencies were not observed in the cropping system with the growth regulator (GR). Durum wheat cultivation at seeding rates of 350 and 450 seeds·m^-2^ contributed to the achievement of the lowest F_max_ value (approx. 115 and approx. 109 N) at nitrogen fertilization dose of 80 kg·ha^-1^ and the highest F_max_ value (approx. 137 and approx. 128N) at nitrogen fertilization dose of 120 kg·ha^-1^. Durum wheat cultivation with the application of GR, at sowing density of 550 seeds·m^-2^, and nitrogen doses of 0, 80, and 120 kg·ha^-1^ allowed producing grain which did not differ significantly in its F_max_ values (approx. 129-136N). Considering the above, it was concluded that the highest maximal compression force (F_max_ of approx. 145 N) was obtained for the grain from WGR system, with sowing density of 350 seeds·m^-2^ and no fertilization with nitrogen (0 kg·ha^-1^) as well as for the grain from the variant with sowing density of 550 seeds·m^-2^ and nitrogen dose of 120 kg·ha^-1^ (approx. 148N) - table 1.

The statistical analysis of the effect of sowing density on durum wheat cultivation in the WGR system demonstrated that this experimental factor caused significant (p<0.01; p<0.05) differences in F_max_ value of the grain at each dose of fertilization, i.e. 0, 80, and 120 kg·ha^-1^. In contrast, in the GR cropping system, this experimental factor had no effect on values of this parameter. The analysis of the GR/WGR effect showed significant (p<0.01; p<0.05) differences in F_max_ of the grain, especially in the variants with lower seeding rates (350 and 450 seeds·m^-2^) and higher fertilization doses (80 and 120 kg·ha^-1^). Therefore, when growing crops in the WGR system more consideration should be given to the sowing density and nitrogen dose, whereas effects of these factors can be neglected in the GR system (tab. 1). According to Spychaj et al. (2011), grain hardness is affected to a greater extent by weather conditions than by chemical plant protection agents used during cultivation. In contrast, Rachoń et al. (2015) demonstrated that the administration of plant protection agents had a positive effect on grain vitreousness, whereas according to Fana et al. (2012) no interaction was observed between fertilization and grain hardness, however this quality parameter positively correlated with flour yield. In turn, Nagamine et al. (2003) showed a significant relationship between flour color and hardness.

### 3.2. Grain color

Results of measurements of durum wheat grain lightness (tab. 2) confirmed results of measurements of the F_max_ value. The highest and similar L* values were determined for the grain from durum wheat grown in the WGR system, at sowing density of 350 seeds·m^-2^ and without nitrogen fertilization (L*=54.35) as well as for the grain of durum wheat fertilized with a nitrogen dose of 80 kg·ha^-1^ (L*=54.16). The lowest L* value (51.36) was demonstrated for the grain from wheat cultivated without the growth regulator, at sowing density of 450 seeds·m^-2^ and nitrogen dose of 120 kg·ha^-1^. As in the case of grain hardness, upon GR use, the L* value of the grain was the highest (53.64) in the cropping with the sowing density of 350 seeds·m^-2^and nitrogen dose of 120 kg·ha^-1^, whereas the lowest one at sowing density of 350 seeds·m^-2^ and nitrogen application of 0 kg·ha^-1^ (51.41). The hardness and lightness of the grain from durum wheat grown in the WGR system were significantly influenced by sowing density, especially at nitrogen fertilization doses of 80 and 120 kg·ha^-1^ (p<0.01; p<0.05) – table 2. The higher L* values of the grain testifies of its lighter color and better usability for semolina and pasta production (Ciolek and Makarska 2004). Results of an experiment conducted by Woźniak (2006) demonstrated intensive fertilization with nitrogen (120 kg·ha^-1^) to significantly increase total ash content in durum wheat grain compared to the minimized fertilization (90 kg·ha^-1^). In turn, according to Sulewska, Koziara and Bojarczuk (2007), nitrogen fertilization decreased ash content and, by this beans, increased the lightness of kernels.

**Table 2.**
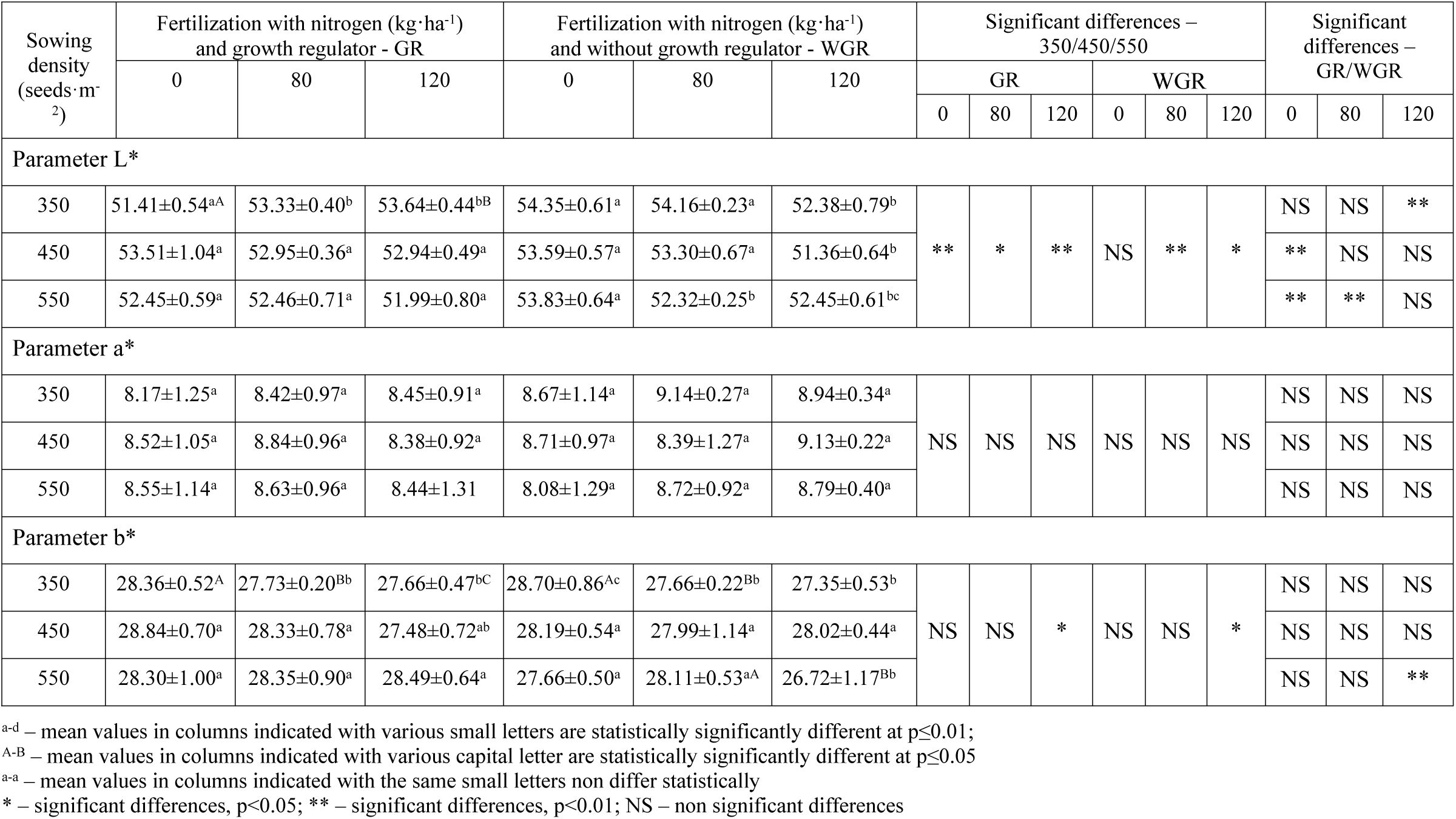
Changes in the values (mean values ± standard deviation) of the color parameters L*, a*, b* durum wheat grain cultivated with the application (GR) and without application (WGR) of the growth regulator with different sowing density and nitrogen fertilization.

The statistical analysis of the effect of nitrogen fertilization dose showed that L* values differed significantly (p<0.01) between the grain from durum wheat grown in the WGR system without fertilization (0N) and that from wheat grown with nitrogen fertilization at doses of 80 and 120 kg·ha^-1^. It was also demonstrated that the sowing density had a significant (p<0.01 and p<0.05) effect on L* values of the grain produced from wheat grown in the GR system in each variant of nitrogen fertilization (0, 80, 120 kg·ha^-1^) as well as on L* values of the grain from durum wheat cultivated in the WGR system with nitrogen fertilization doses of 80 and 120 kg·ha^-1^ (p<0.05). The analysis of GR/WGR effect showed that this factor caused the greatest differences in L* values of the grain from wheat grown at 0 kg·ha^-1^ and sowing density of 450 and 550 seeds·m^-2^, at 80 kg·ha^-1^ /550 seeds·m^-2^, and at 120 kg·ha^-1^ /350 seeds·m^-2^ (tab. 2).

Results of measurements of grain redness (a*) demonstrated that its values were similar (8.17 – 9.13) and were not significantly affected by any of the factors tested (WGR/GR, 350/450/550; 0/80/120 kg·ha^-1^) – table 2.

Results of measurements of grain yellowness (b*), being indicative of the content of carotenoid pigments (Hentschel et al. 2002; Ciołek, and Makarska 2004; Colasuonno et al. 2019), demonstrated its values to range from 26.72 to 28.84 regardless of the analyzed variant and experimental factor. The highest saturation with yellow color (b*≈29) was determined in the grain from durum wheat grown at GR/0N/450 seeds·m^-2^. The b* values obtained in the study (approx. 27.3 or higher) point to the good quality of the grain (Colasuonno et al. 2019; Zhang and Dubcovsky 2008). Durum grain with a higher content of yellow pigments is characterized by superior qualitative composition of gluten proteins and it has lighter and thinner hulls. In turn, the achieved lowest b* values (GR/450: 27.48; WGR/550: 26.72) and the observed significant (p<0.01) effect of WGR/GR and seeding rate (350/450/550) on b* values of the grain from durum wheat fertilized with a nitrogen dose of 120 kg·ha^-1^ (tab. 2) are indicative of the negative effect of fertilization with nitrogen on the content of carotenoid pigments (Sulewska, Koziara and Bojarczuk 2007). The levels of lightness (L*) and redness (a*) values of analyzed durum grain were relatively similar to these obtained for common wheat and spelt grain, whereas values of durum grain yellowness (b*) were 1.5x higher than in the case of mentioned two wheat species (Zuk-Golaszewska et al. 2018).

### 3.3. Flour extraction rate (FER)

Being the basic tool used to monitor and control the milling process as well as its technological and economic outcomes in the milling industry, the flour extraction rate (FER) was described as a quality indicator which allows establishing the optimal conditions of crop cultivation (Zuk-Golaszewska et al. 2018). The yield of semolina declines as kernels become thinner because the proportion of endosperm concomitantly decreases. This is largely influenced by environmental conditions during the growing season and during harvest (Owens 2001).

Results of determination of the amount of flour produced from grain unit demonstrated a higher extraction rate of the flour made of the grain from durum wheat grown in the WGR system (68.69-59.38%), compared to the GR system (64.00 – 58.58%), however the highest FER value was obtained in the variant with 0N and 350 seeds m^-2^ (68.96%) which allowed producing the hardest grain (tab.1). In addition, the sowing density factor (350/450/550) in the WGR system, caused significant (p<0.01, p<0.05) differences in FER, regardless of nitrogen fertilization dose. The lower FER value of flour from the grain of wheat cultivated in the GR system was also significantly (p<0.01) affected by sowing density at 80N fertilization dose. The highest FER values, being significantly different from these obtained at WGR/80N, were achieved during wheat cultivation at 80N fertilization and sowing densities of 350 seeds·m^-2^ (64.00%) and 450 seeds m^-2^ (63.92%) – table 3. Obtained FER values were comparable or even higher than domestic standards for this parameter, which were: 66.1% – spring common wheat cv. Tybalt; 61,7% – spring durum wheat cv. SMH87 (Polish National List of the Research Center for Cultivar Testing. 2011).

**Table 3.** Changes in the extraction rate and particle size (mean values ± standard deviation) of flour from durum wheat grain depending on sowing density, growth regulator application and nitrogen fertilization.

| Sowing density<br>(seeds·m<sup>-2</sup>) |  | FER (%) |  |  |  |  |  | FPS (%) |  |  |  |  |  |
| --- | --- | --- | --- | --- | --- | --- | --- | --- | --- | --- | --- | --- | --- |
|  |  | Nitrogen fertilization (kg·ha<sup>-1</sup>) |  |  | Sig. diff. –<br>350/450/550 |  |  | Nitrogen fertilization (kg·ha<sup>-1</sup>) |  |  | Sig. diff. –<br>350/450/550 |  |  |
|  |  | 0 | 80 | 120 | 0 | 80 | 120 | 0 | 80 | 120 | 0 | 80 | 120 |
| Without the application growth regulator (WGR) |  |  |  |  |  |  |  |  |  |  |  |  |  |
| 350 |  | 68.96±1.10<sup>a</sup> | 59.91±2.18<sup>a</sup> | 60.15±0.89<sup>a</sup> | ** | ** | * | 67.60±0.75<sup>a</sup> | 97.11±0.07<sup>a</sup> | 95.94±0.76<sup>a</sup> | ** | ** | ** |
| 450 |  | 67.52±1.09<sup>a</sup> | 60.45±0.44<sup>a</sup> | 62.02±0.92<sup>aA</sup> |  |  |  | 92.83±1.44<sup>b</sup> | 98.36±0.41<sup>bB</sup> | 99.57±0.15<sup>b</sup> |  |  |  |
| 550 |  | 59.38±1.07<sup>b</sup> | 64.31±1.11<sup>b</sup> | 59.67±0.98<sup>Ba</sup> |  |  |  | 95.00±0.20<sup>b</sup> | 97.64±0.12<sup>Cc</sup> | 97.09±0.16<sup>ca</sup> |  |  |  |
| With the application growth regulator (GR) |  |  |  |  |  |  |  |  |  |  |  |  |  |
| 350 |  | 62.05±1.65<sup>a</sup> | 64.00±0.40<sup>a</sup> | 60.94±0.23<sup>a</sup> | NS | ** | NS | 96.31±0.66<sup>a</sup> | 94.25±0.93<sup>a</sup> | 99.77±0.07<sup>a</sup> | ** | ** | NS |
| 450 |  | 62.99±0.71<sup>a</sup> | 63.92±0.89<sup>a</sup> | 58.58±1.70<sup>a</sup> |  |  |  | 91.03±0.84<sup>b</sup> | 93.62±0.11<sup>a</sup> | 99.37±0.06<sup>aB</sup> |  |  |  |
| 550 |  | 61.28±1.51<sup>a</sup> | 61.22±0.63<sup>b</sup> | 59.82±1.40<sup>a</sup> |  |  |  | 97.11±0.57<sup>ca</sup> | 97.97±0.09<sup>b</sup> | 99.40±0.38<sup>Ca</sup> |  |  |  |
| Significant<br>differences<br>–<br>GR/WGR | 350 | ** | ** | NS | - | - | - | ** | ** | ** | - | - | - |
|  | 450 | ** | ** | * |  |  |  | NS | ** | NS |  |  |  |
|  | 550 | NS | * | NS |  |  |  | ** | * | ** |  |  |  |

FER - flour extraction rate; FPS – flour particle size.
<sup>a-d</sup> – mean values in columns indicated with various small letters are statistically significantly different at $p < 0.01$ ;
<sup>A-B</sup> – mean values in columns indicated with various capital letter are statistically significantly different at $p < 0.05$
<sup>a-a</sup> – mean values in columns indicated with the same small letters non differ statistically
\* – significant differences, $p < 0.05$ ; \*\* – significant differences, $p < 0.01$ ; NS – no significant differences

### 3.4. Flour particle size (FPS)

The particle size distribution of flour (FPS) is another quality indicator which affects its storage stability and technological usability (e.g. water absorption rate or dough formation time). Pasta producers are fully aware of the impact the particle size of durum wheat semolina or flour has on the pasts quality. However, detailed requirements regarding this quality indicator differ significantly among countries. Due to the great advance in novel techniques and technologies employed in the pasta making process, the requirements set for pasta raw materials regarding particle size distribution have changed remarkably in recent years. The demand for coarse pasta flours has decreased, while that for semolina and durum flours with finer but uniform granulation has increased (Jurga 2007). For years, pasta has been produced with semolina having particle sizes of 125 to 630 μm, whereas today the most common and suitable raw material is that with finer particles being smaller than 400 μm (Polish National List of the Research Center for Cultivar Testing 2011). This allows shortening kneading time and achieving pasta dough with a more homogenous structure.

Results of FPS measurements demonstrate that flour from durum wheat grown in the WGR system was characterized by a wider range of values of this parameter (67.6 – 99.57%) and, that the lowest value was determined in the flour made of the grain from wheat grown without nitrogen fertilization at sowing density of 350 seeds m^-2^. In turn, in the GR system, the range of FPS values was narrower, but the values achieved were higher, i.e. 91.03 – 99.77% (table 3).

The analysis of the effect of WGR/GR factor showed significant (p<0.01, p<0.05) differences in the FPS distribution of flour. In the cropping variants: 0/120 kg·ha^-1^ – 350 and 550 seeds m^-2^, the FPS value of flour from WGR grain was lower compared to that of flour from GR grain. In contrast, the FPS values were significantly (p<0.05) higher in the case of flours from WGR grain and 80 kg·ha^-1^ variant (at all sowing densities) – table 3. Obtained FPS values were also comparable or higher than domestic standards for this parameter, which were: 91.6% – spring common wheat cv. Tybalt; 82.2% – spring durum wheat cv. SMH87 (Polish National List of the Research Center for Cultivar Testing 2011).

### 3.5. Flour color

The highest values of L* color parameter of flour (91.93-91.81) of durum grain produced in WGR system were obtained for the variants WGR/80 kg·ha^-1^ /350-450 seeds·m^-2^ and WGR/0N/550 seeds·m^-2^. In case of GR system, a highest range of L* values (92.43-91.74) was reported for the flours made of the grain from the variants GR/0 kg·ha^-1^/350-550 seeds·m^-2^. In flours from the other variants, L* values ranged from 90.48 to 91.12. The analysis of the impact of agrotechnical factors demonstrated a significant (p<0.01; p<0.05) effect of GR/WGR on the lightness of flours from all cropping variants, except for the variant 120 kg·ha^-1^/450 seeds·m^-2^. The sowing density factor caused significant differences in L* values of flours made of the grain from the variant GR/0; 120 kg·ha^-1^ (p<0.01; p<0.05) and from the variant WGR/0; 80; 120 kg·ha^-1^ (p<0.01) – table 4.

**Table 4.** Changes in the values (mean values ± standard deviation) of the color parameters L*, a*, b* of flour from grain of durum wheat cultivated with the application (GR) and without application (WGR) of the growth regulator with different sowing density and nitrogen fertilization.

| Sowing density<br>[seeds·m<sup>-2</sup>] | Fertilization with nitrogen [kg·ha<sup>-1</sup>]<br>and growth regulator - GR |  |  | Fertilization with nitrogen [kg·ha<sup>-1</sup>]<br>and without growth regulator - WGR |  |  | Significant differences –<br>350/450/550 |  |  |  |  |  | Significant<br>differences –<br>GR/WGR |  |  |
| --- | --- | --- | --- | --- | --- | --- | --- | --- | --- | --- | --- | --- | --- | --- | --- |
|  | 0 | 80 | 120 | 0 | 80 | 120 | GR |  |  | WGR |  |  |  |  |  |
|  |  |  |  |  |  |  | 0 | 80 | 120 | 0 | 80 | 120 | 0 | 80 | 120 |
| Parameter L* |  |  |  |  |  |  |  |  |  |  |  |  |  |  |  |
| 350 | 91.74±0.20<sup>a</sup> | 90.90±0.16<sup>aB</sup> | 90.60±0.25<sup>aC</sup> | 90.85±0.29<sup>a</sup> | 91.80±0.16<sup>a</sup> | 91.05±0.30<sup>a</sup> | ** | NS | * | ** | ** | ** | ** | ** | * |
| 450 | 91.78±0.23<sup>a</sup> | 91.12±0.30<sup>aA</sup> | 90.50±0.32<sup>aA</sup> | 90.96±0.67<sup>aA</sup> | 91.93±0.30<sup>a</sup> | 90.48±0.20<sup>b</sup> |  |  |  |  |  |  | * | ** | NS |
| 550 | 92.43±0.43<sup>b</sup> | 90.77±0.23<sup>B</sup> | 90.90±0.16<sup>B</sup> | 91.81±0.34<sup>Bb</sup> | 91.07±0.24<sup>b</sup> | 90.50±0.17<sup>b</sup> |  |  |  |  |  |  | * | * | ** |
| Parameter a* |  |  |  |  |  |  |  |  |  |  |  |  |  |  |  |
| 350 | 0.41±0.08<sup>aB</sup> | 0.62±0.13<sup>a</sup> | 0.45±0.09<sup>a</sup> | 0.66±0.09<sup>a</sup> | 0.58±0.03<sup>a</sup> | 0.51±0.06<sup>a</sup> | NS | ** | NS | ** | NS | ** | ** | NS | NS |
| 450 | 0.50±0.10<sup>aA</sup> | 0.56±0.05<sup>a</sup> | 0.42±0.07<sup>a</sup> | 0.63±0.03<sup>a</sup> | 0.59±0.07<sup>a</sup> | 0.55±0.06<sup>a</sup> |  |  |  |  |  |  | * | NS | ** |
| 550 | 0.40±0.05<sup>B</sup> | 0.45±0.02<sup>b</sup> | 0.42±0.04<sup>a</sup> | 0.43±0.08<sup>b</sup> | 0.68±0.12<sup>b</sup> | 0.65±0.04<sup>b</sup> |  |  |  |  |  |  | NS | ** | ** |
| Parameter b* |  |  |  |  |  |  |  |  |  |  |  |  |  |  |  |
| 350 | 18.22±0.52<sup>aB</sup> | 19.30±0.89<sup>a</sup> | 19.14±0.31<sup>a</sup> | 17.82±0.24<sup>a</sup> | 19.22±0.29<sup>aC</sup> | 19.43±0.13<sup>a</sup> | NS | NS | * | ** | ** | NS | NS | NS | NS |
| 450 | 18.55±0.18<sup>aA</sup> | 19.54±0.21<sup>a</sup> | 19.52±0.35<sup>a</sup> | 18.60±0.40<sup>b</sup> | 19.69±0.10<sup>bA</sup> | 19.41±0.19<sup>a</sup> |  |  |  |  |  |  | NS | NS | NS |
| 550 | 18.12±0.30<sup>B</sup> | 19.51±0.41<sup>a</sup> | 19.73±0.40<sup>a</sup> | 18.21±0.33<sup>b</sup> | 19.52±0.09<sup>B</sup> | 19.66±0.51<sup>b</sup> |  |  |  |  |  |  | NS | NS | NS |

a-d – mean values in columns indicated with various small letters are statistically significantly different at p<0.01;
A-B – mean values in columns indicated with various capital letter are statistically significantly different at p<0.05
a-a – mean values in columns indicated with the same small letters non differ statistically
\* – significant differences, p<0.05; \*\* – significant differences, p<0.01; NS – non significant difference

Results of redness measurements demonstrated that a* values of the flour made of the grain from the WGR system were higher (0.68-0.43) compared to a* values of flour from wheat grain from the GR system (0.62-0.40). The statistical analysis of the impact of agrotechnical factors demonstrated a significant (p<0.01; p<0.05) effect of the GR/WGR factor on a* value of the flours made of the grain produced from variants: 0 kg·ha^-1^/350; 450 seeds·m^-2^; 80 kg·ha^-1^/550 seeds·m^-2^, and 120 kg·ha^-1^/450 and 550 seeds·m^-2^. Sowing density caused significant (p<0.01) differences in a* values of flours produced from the grain from variants: WGR/0; 120 kg·ha^-1^ and GR/80 kg·ha^-1^ (table 4).

The statistical analysis of the impact of agrotechnical factors demonstrated that only sowing density caused significant differences the yellowness (b*) of flour made of the grain produced from variants: GR/120 kg·ha^-1^ (p<0.05) and WGR/0; 80 kg·ha^-1^ (p<0.01) – table 4. The lack of an explicit effect of the agrotechnical factors on the yellowness of flour, like in the case of analyzed parameter b* of grain color, may be due to the complex nature of the yellow pigment in semolina/ flour from durum wheat grain, whose carotenoid fraction accounted for only 30-50% of the yellow pigment quantities. There are still compounds in durum wheat not yet identified that contribute considerably to the yellow color of the flour (Hentschel et al. 2002; Colasuonno et al. 2019). The level of lightness L* values of analyzed durum wheat flour was relatively similar to this obtained for flours of common wheat and spelt grain, whereas values of durum flour redness a* and yellowness b* were twice higher than in the case of mentioned two wheat species (Zuk-Golaszewska et al. 2018).

### 3.6. Correlations

To determine the effect of agrotechnical factors (seeding density, nitrogen fertilization, retardant application) on the analyzed parameters of grain quality and flour obtained from it, an analysis of correlations between these variables was performed.

Analysis of the influence of sowing density and nitrogen fertilization grain cultivated without the application of growth regulator (WGR) showed that nitrogen fertilization had a significant (p <0.01) and a strong effect on the color parameters L* (r = -0.665) and b* (r = -0.488) of the grain, and these correlations were negative (with increasing nitrogen dose, parameter values decreased). The sowing density was characterized by a smaller effect on the lightness of the grain (r = -0.286). The opposite relationship was observed for grain cultivated with the application of growth regulator (GR). Greater and negative correlation (r = -0.676, p <0.01) was obtained between sowing density of grain and its lightness (L*), while the effect of nitrogen fertilization on the hardness and its color parameters (L*, b*) was characterized by a significant, but of a smaller strength, correlation (p <0.05) - table 5.

**Table 5.**
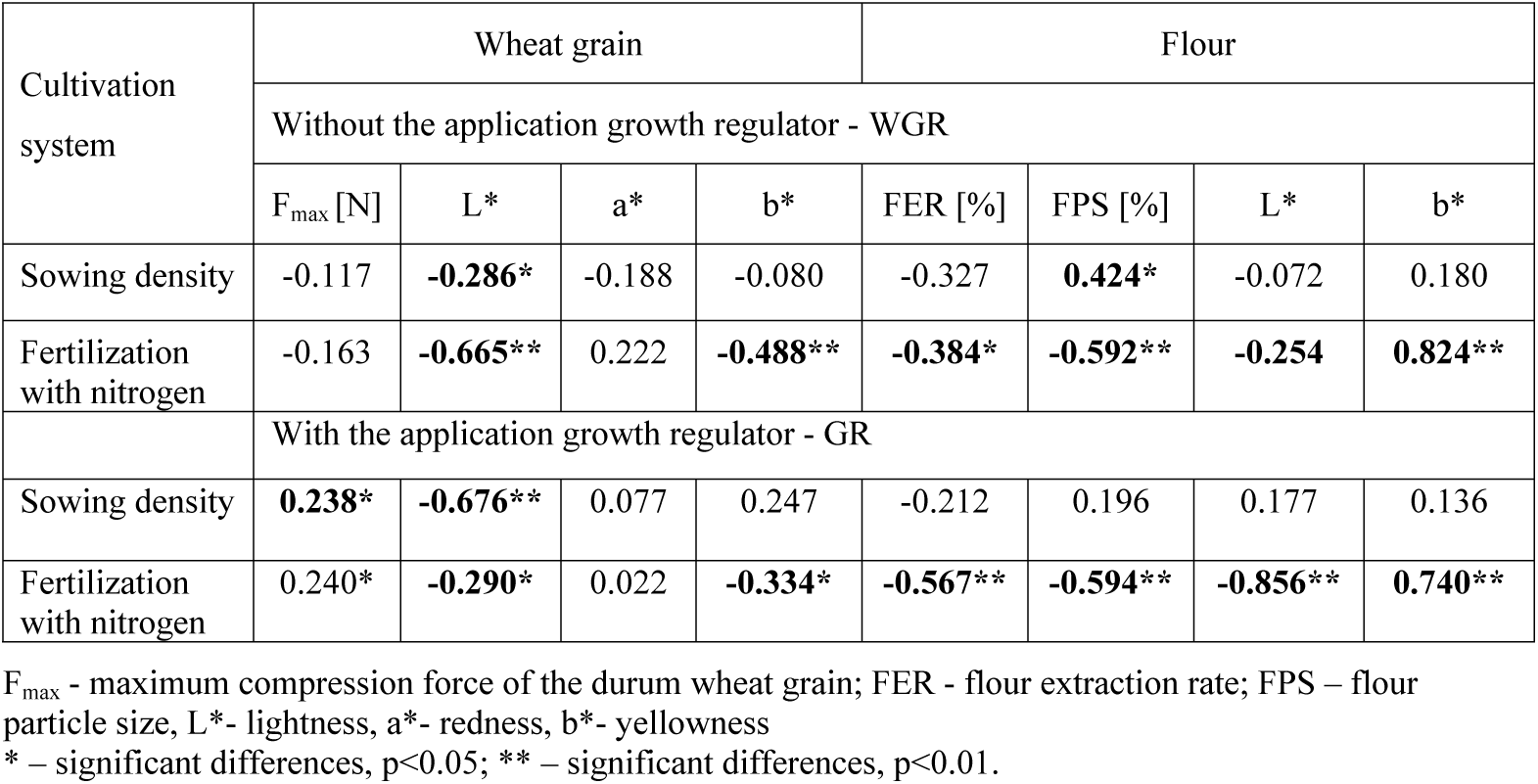
Correlation coefficients between agrotechnical factors (sowing density, fertilization with nitrogen, WGR / GR) and selected qualitative parameters durum wheat grain and flour made of it.

| Cultivation system | Wheat grain |  |  |  | Flour |  |  |  |
| --- | --- | --- | --- | --- | --- | --- | --- | --- |
|  | Without the application growth regulator - WGR |  |  |  |  |  |  |  |
|  | F<sub>max</sub> [N] | L* | a* | b* | FER [%] | FPS [%] | L* | b* |
| Sowing density | -0.117 | **-0.286\*** | -0.188 | -0.080 | -0.327 | **0.424\*** | -0.072 | 0.180 |
| Fertilization with nitrogen | -0.163 | **-0.665\*\*** | 0.222 | **-0.488\*\*** | **-0.384\*** | **-0.592\*\*** | **-0.254** | **0.824\*\*** |
|  | With the application growth regulator - GR |  |  |  |  |  |  |  |
| Sowing density | **0.238\*** | **-0.676\*\*** | 0.077 | 0.247 | -0.212 | 0.196 | 0.177 | 0.136 |
| Fertilization with nitrogen | 0.240* | **-0.290\*** | 0.022 | **-0.334\*** | **-0.567\*\*** | **-0.594\*\*** | **-0.856\*\*** | **0.740\*\*** |

F<sub>max</sub> - maximum compression force of the durum wheat grain; FER - flour extraction rate; FPS – flour particle size, L\*- lightness, a\*- redness, b\*- yellowness \* – significant differences, $p < 0.05$ ; \*\* – significant differences, $p < 0.01$ .

In the case of flour obtained from grain cultivated without the application of growth regulator (WGR), similar relationships were observed as for the grain itself. The most significant (p <0.01) and strong effect of nitrogen fertilization was obtained for FPS (r = -0.592) and yellowness b* (r = 0.824), and a slightly lower strength - for its FER (r = -0.384). However, the grain sowing density also significantly affected (p <0.05) on the FPS. In the case of flour obtained from grain cultivated with the application of growth regulator (GR), significant effect (p <0.01) and high strength (r = (−0.856) - (0.740)) on the all tested technological parameters had nitrogen fertilization - table 5.

To determine the effect of application of GR during cultivation, the Mann-Witney’s test was carried out for 2 independent groups. The obtained results did not allow the rejection of the null hypothesis (the means of the compared groups are equal), which means that the impact of sowing density and nitrogen fertilization on both groups is similar - statistically insignificant.

## 4. Conclusions

1. Grain of the highest hardness was produced from durum wheat grown without the growth regulator (WGR), at the lowest sowing density (350 seeds·m^-2^) and nitrogen fertilization dose of 80 kg·ha^-1^.
2. The highest values of lightness (L*) and yellowness (b*) were determined in the grain of wheat cultivated without additional agrotechnical measures (growth regulator and nitrogen fertilization). The lightness (L*) of the grain produced from wheat cultivated with the support of the growth regulator was negatively affected by increasing the sowing density to 550 seeds·m^-2^ and nitrogen fertilization dose to 120 kg·ha^-1^.
3. Study results, supported by correlation analysis, indicated that high-quality grain with desired flour quality parameters (FER, FPS, L*) can be produced from spring durum wheat grown without the growth regulator and at medium doses of nitrogen fertilization.
4. Obtaining good quality technological parameters of durum grain and flour in case of application medium doses nitrogen and without the growth regulator can reduce costs of durum wheat production and contamination of the natural environment.

